# Porechop_ABI: discovering unknown adapters in ONT sequencing reads for downstream trimming

**DOI:** 10.1101/2022.07.07.499093

**Authors:** Quentin Bonenfant, Laurent Noé, Hélène Touzet

## Abstract

**Motivation:** Oxford Nanopore Technologies (ONT) sequencing has become very popular over the past few years and offers a cost-effective solution for many genomic and transcriptomic projects. One distinctive feature of the technology is that the protocol includes ligation of adapters to both ends of each fragment. Those adapters should then be removed before downstream analyses, either during the basecalling step or by explicit trimming. This basic task may be tricky when the definition of the adapter sequence is not well-documented.

**Results:** We have developed a new method to scan a set of ONT reads to see if it contains adapters, without any prior knowledge on the sequence of the potential adapters, and then trim out those adapters. The algorithm is based on approximate *k*-mers and is able to discover adapter sequences based on their frequency alone. The method was successfully tested on a variety of ONT datasets with different flowcells, sequencing kits and basecallers.

**Availability:** The resulting software, named Porechop_ABI, is open-source and is available at https://github.com/bonsai-team/Porechop_ABI.

## 1 Introduction

ONT is a versatile sequencing technology that produces long reads and has many applications, including *de novo* genome sequencing, metagenomics [5], analysis of structural variants [7] and transcriptome sequencing [12, 8]. One common feature to all sequencing kits and flowcells is that library preparation includes ligation of adapter sequences to both ends of DNA, cDNA or RNA fragments. These adapters facilitate strand capture and loading of a processive enzyme.

Since the adapters are sequenced with the fragment, this implies that each resulting read may contain either full-length or partial adapters due to incomplete sequencing.

In this context, read trimming is a mandatory pre-processing step to remove these artificial sequences from raw reads and avoid unexpected interference in downstream analyses. Tools such as Porechop, that finds and removes adapters, were designed to perform this task efficiently [10]. The main limitation however is that such tools rely on a static database of known adapter sequences, which is a critical issue when the adapter sequences used are not known, or when there is no information about the fact that the reads have already been trimmed out or not. Additionally, Porechop and its database are no longer maintained since october 2018. More recently, ONT released the Guppy toolkit that contains several basecalling and post-processing algorithms, including adapter trimming. But this toolkit can be seen as a black box with no control on the output. Moreover, it cannot be applied to previously published public datasets when the FAST5 files are no longer available.

To solve these problems, we have developed a new software to automatically infer adapter sequences from raw reads alone, without any external knowledge or database. This algorithm determines whether the reads contain adapters, and if so what the content of these adapters is. It uses technics coming from string algorithms, with approximate *k*-mer, full text indexing and assembly graphs.

The method is available as an extension of the existing Porechop tool, and the resulting software is named Porechop_ABI. This new tool is proving to be useful to clean untrimmed reads for which the adapter sequences are not documented, to check whether a dataset has been trimmed or not. It is even able to find leftover adapters in datasets that been previously processed with Guppy with trimming mode activated, or to deal with datasets with several distinct adapters.

## 2 Algorithm

The goal is to design a computational method that is able to infer, or to guess, the adapter sequences from a set of untrimmed reads. The starting point of the method is that adapters are expected to be found mainly at each extremity on untrimmed reads and are over-represented sequences that could be distinguished from the biological content. To work properly, the method should fulfill several additional constraints: it should be tolerant of sequencing errors; it should scale to large datasets; it should deal with adapters of varying length (from 16nt to more than 30nt); it should accomodate to the presence of several distinct adapters in the data set.

For that, we have developed a new algorithm that is based on four main steps :

i. Reads sampling: Select 10 independent samples of 40,000 reads from the dataset, then for each reads of the samples select start and end regions of length 100 nt
ii. Approximate *k*-mer counting: Find and count *k*-mers that are over-represented throughout the start (resp. end) region. This search allows for edit errors (insertions, deletions and substitutions)
iii. Adapter construction: Reconstruct the start (resp. end) adapter sequence by assembling *k*-mers using an assembly graph based on most represented *k*-mers
iv. Consensus between samples: Align and compare the start (resp. end) adapters found for each of the 10 samples, and build a consensus sequence. When the sequences are not fully compatible, when there is no single consensus sequence, the method outputs several adapter sequences associated to a support score, that corresponds to the proportion of samples containing the adapter.

We detail each of these steps in Subsections 2.1, 2.2 and 2.3, which consitutes the *core module*. This core module is then used in a sampling and consensus approach, that is described in Subsection 2.4. Finally, the choice of the parameters is discussed in Subsection 2.5. The overal organization of the tool is given in Figure.

### 2.1 Selection of the most frequent *k*-mers

#### 2.1.1 Overal principle

The starting point of the algorithm is to search for frequent *k*-mers appearing at the beginning (they are included in the region [1, 100] for the start adapter) or at the end (they are included in the region [−99, −0] for the end adapter) of the reads. This step is motivated by a simple observation. In a raw dataset with untrimmed adapters, most of the reads include the adapters in these regions, whereas the content of the biological sequence in between is more variable. So the *k*-mers composing the adapter sequences should be more frequent than the other *k*-mers occurring in the dataset.

For each of the two regions separately, we search for the 500 most frequent *k*-mers. Authorized values of *k* for *k*-mer lengths vary from 2 to 32. In practice, recommended values range between 16 and 22, and the default value is *k* = 16 which is a compromise between the length of adapter sequences and the error sequencing rate (see Subsection 2.5).

#### 2.1.2 Low complexity *k*-mers

During the *k*-mer count, we filter out *low complexity k-mers*. Indeed, such *k*-mers can have high counts despite not being part of the adapter. Common examples are homopolymers or poly-A tail of mRNA. To detect low complexity *k*-mers, we use a complexity score inspired by the DUST score [4]. The difference is that we use dimers instead of triplets, because of the shorter length of the sequences.

The definition of the score is as follows. Let *a* be a *k*-mer (*k* > 2) and let ℛ be the set of all 16 possible dimers on the alphabet {*A, T, C, G*}. We define *c*_*d*_(*a*) to be the number of occurrences of a dimer *d* ∈ ℛ in the *k*-mer *a*. The *low complexity score* of *a*, denoted *Lc*(*a*), is :

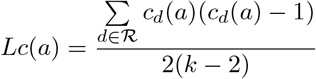

The higher the *Lc* score, the lower the complexity of the *k*-mer. We define a a score threshold above which the *k*-mer is rejected. This score depends on the value *k*. For *k* = 16, it equals *lc*_16_ = 1.0, which amounts to discard approximately 2.7% of all possible 16-mers. For other values of *k*, we automatically readjust the score threshold using a regression wrt *k* = 16:

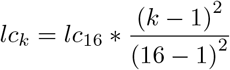

### 2.2 Computation of the 2-errors frequency

The first step allowed to identify *exact* frequent *k*-mers. Because of the high error rate of ONT reads, we also expect to have sequencing errors in the adapters. This means that some erroneous *k*-mers will be found inside the 500 selected *k*-mers, and that some actual *k*-mers may have a higher frequency in the initial data than that observed frequency in the sequencing data.

To take this phenomenon into account, we perform a second count and compute for each *k*-mer its number of occurrences with *up to two errors* (substitutions, insertions or deletions). We call it the 2*-error frequency* of the *k*-mer. To do that, we re-scan the start and end regions of the sample, and compute for each of the top 500 *k*-mers previously found its number of approximate occurences. This search is performed efficiently by the use of the Optimal Search Scheme algorithm [3] implemented in the SeqAn library [6].

### 2.3 Reconstruction of the adapter sequence

The last step of the core module aims at reconstructing the adapter sequences from the *k*-mers. It is based on an assembly graph. The nodes of the graph are the 500 most frequent *k*-mers weighted by their 2-errors frequency. There is an arc between two nodes when these nodes have an overlap of length *k* − 1 (as in a De Bruijn graph). The goal is to find a path in the graph that corresponds to the original adapter sequence.

#### 2.3.1 Path construction in the graph

Our approach is to search for the *heaviest path*, where the weight of a path is defined as the sum of the weights of the nodes composing the path. This is done by dynamic programming, using the algorithm available in the NetworkX library (https://networkx.org/).

#### 2.3.2 Boundaries of the adapter sequence within the path

The construction of the path tends to make the resulting path longer than the actual adapter, because it keeps incorporating *k*-mers as long as *k*-mer nodes are available in the graph. It means that the actual adapter sequence is a substring of the path.

So there is a need to devise a specific method that is able to limit the impact of the outlier *k*-mers with very high or low counts relative to their neighbouring *k*-mers.

Let *n*_1_, …, *n*_*ℓ*_ be the nodes composing the path for the start adapter sorted from left to right. We smooth the distribution of 2-errors frequencies by using a sliding window median approach. For a given position *i* in the path, we consider the *w* preceding nodes *n*_*i*−*w*_, … *n*_*i*−1_ and the *w* following nodes *n*_*i*+1_, …, *n*_*i*+*w*_. The value *f*_*i*_ is defined as the median values of the 2*w* + 1 2-errors frequencies. We then search for consecutive positions in the path with higher *contrast*. It means that from the list *f*_1_, …, *f*_*ℓ*_, we select the largest index *j* such that *f*_*j*_ − *f*_*j*+1_ is greater than a given threshold. This threshold corresponds to a drop in the curve. It is defined as the median value of *f*_*i*_ − *f*_*i*+1_ plus 7.5% of the maximum count. The length of the window and the 7.5% threshold are determined empirically. A window of size 7 (corresponding to *w* = 3) yields the best results, with some variations on very small raw adapters (≤ 22 bases, or if too close to the *k*-mer size). Selecting the largest value for *j* corresponds to the longest prefix, and allows to capture series of consecutive adapters. All this process is illustrated in Figure 2. If no such threshold exists, we keep the entire sequence: there is no need to readjust the boundaries.

**Figure 1:**
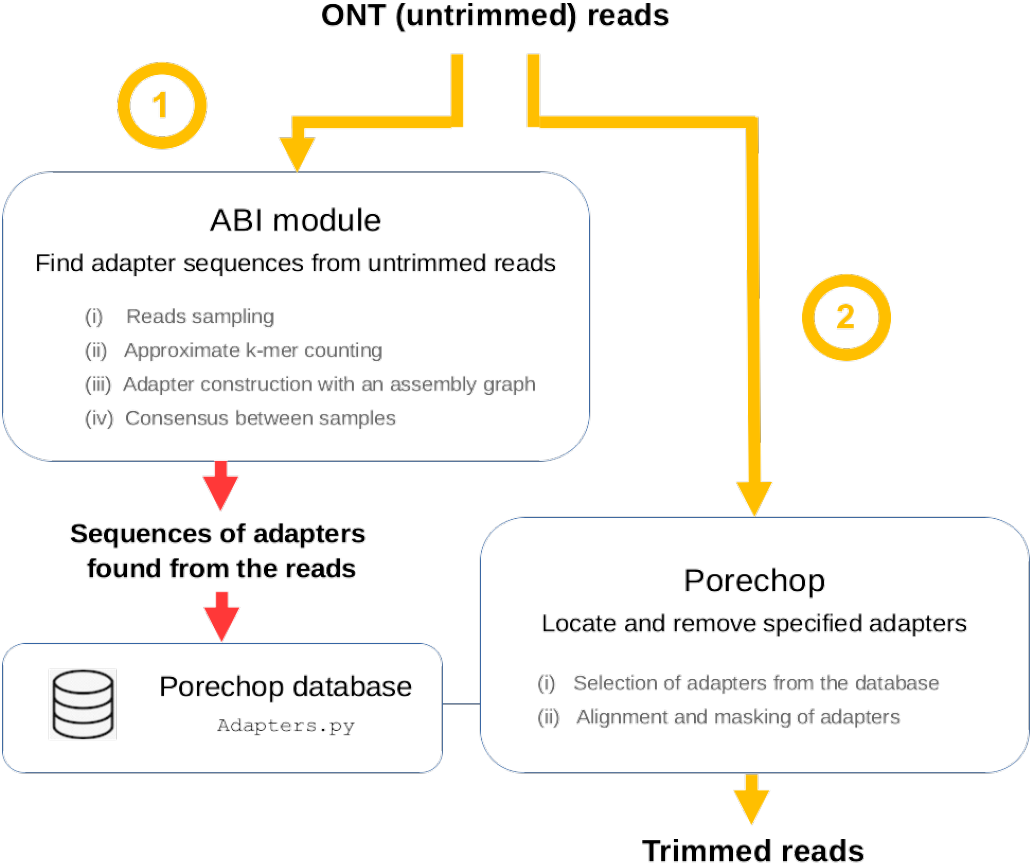
Organization of Porechop_ABI. (1) The tool takes as input a set of reads. When those reads are untrimmed, the ABI module is able to determine the sequences of the adapters that have been used in the sequencing protocol. (2) Those sequences are then ready to be used by Porechop for downstream trimming. If the reads are already trimmed out, no adapter sequence is returned by the ABI module.

**Figure 2:**
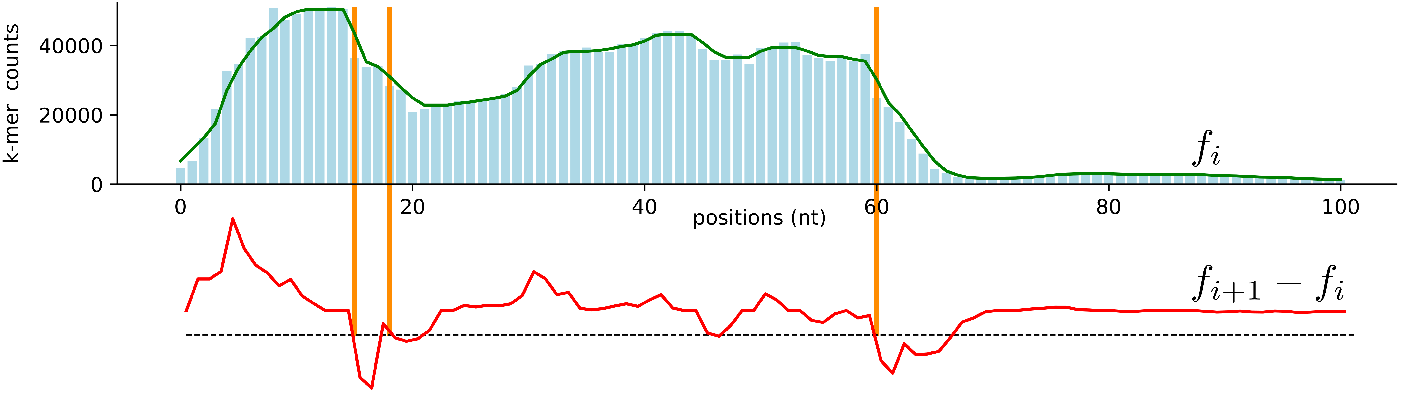
Boundaries of the adapter sequence within the path

Similarly, we consider the path *e*_1_, …, *e*_*m*_ obtained for the end adapter, construct the smoothed distribution of 2-errors frequency and search for the longest suffix *e*_*i*_, … *e*_*m*_, which is characterised by the smallest position *i* such that *e*_*i*+1_ − *e*_*i*_ is greater than the threshold.

#### 2.3.3 Low frequency warning

The general principle of the algorithm, through the different steps, is to exploit the frequency differences between *k*-mers of the adapter and the other *k*-mers present in the reads. In case the most frequent *k*-mers do not show a clear over-representation, the result of the algorithm is questionable, because the signal is not reliable. This may happen when the input sequences are already trimmed, for example. Such case is easily detected. When the 2-errors frequency of the *k*-mer with the highest 2-errors frequency is smaller than 10% of the total number of sequences of the sample (by default 40,000), the program outputs a *low frequency warning* message, meaning that the adapter may be unstable or that the dataset is not suitable for trimming.

### 2.4 Sampling and consensus

Subsections 2.1, 2.2 and 2.3 constitute what we call the *core module* of Porechop_ABI. We do not run this core module on the whole set of reads, because it would be time-consuming. Instead, we work with random samples of the set of reads. This sampling strategy is described in Figure 3. It has several benefits: this speeds up the computation, this allows to capture diversity between the samples and this allows to improve the prediction through the construction of consensus sequences between samples.

**Figure 3:**
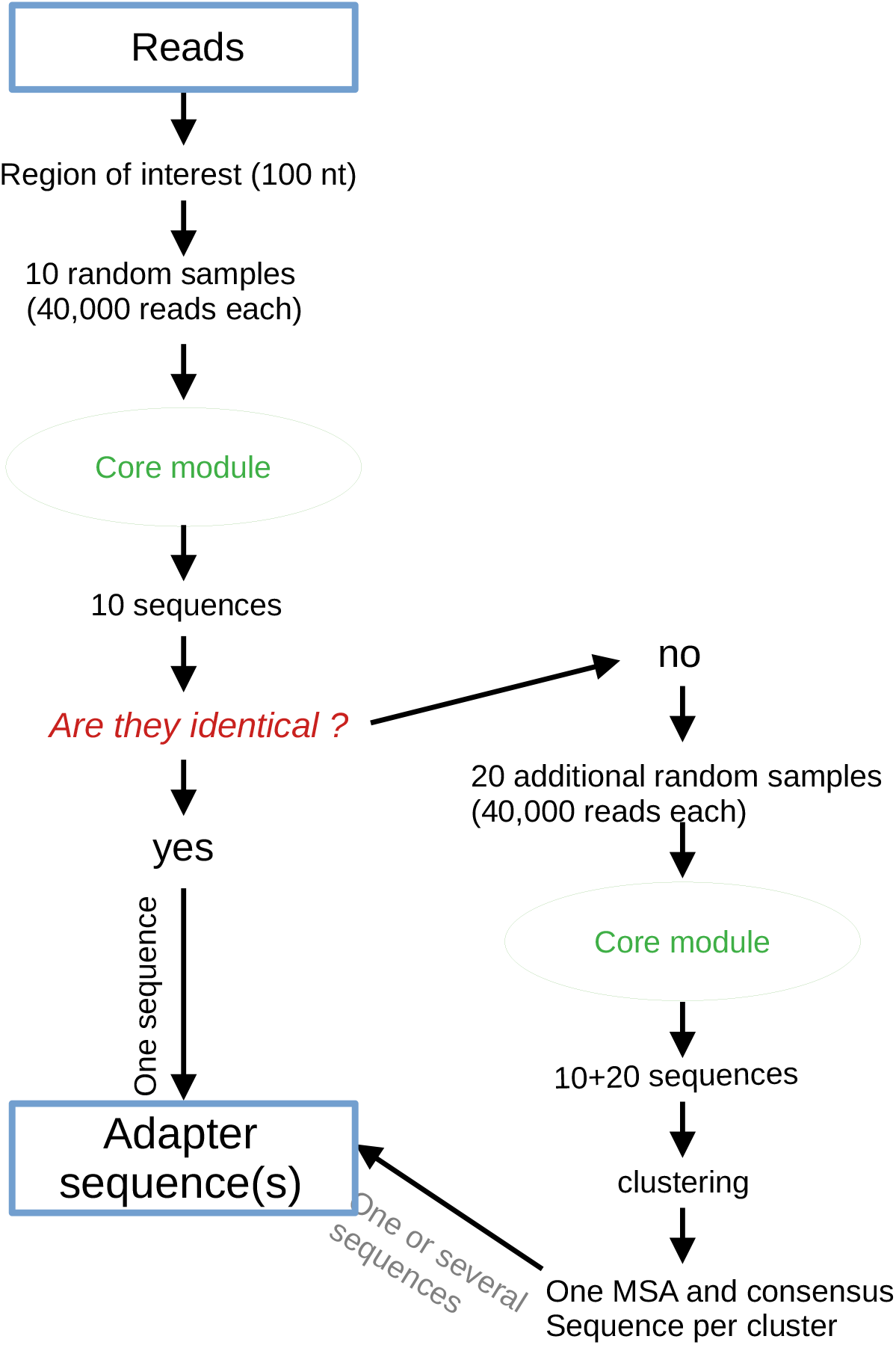
Sampling strategy with the core module

#### 2.4.1 Sampling

We first randomly pick up 10 datasets of 40,000 reads with length at least 200 nt, and compute one putative sequence for each of these datasets using the algorithm described previously in Subsections 2.1, 2.2 and 2.3. If the 10 sequences found for each sample are identical, we stop and output this sequence: this is the adapter sequence. If not, we pick up 20 more random samples of 40,000 reads, and compute a candidate sequence for each of these datasets. This gives a total number of 30 sequences, for which we build a consensus sequence.

#### 2.4.2 Building consensus sequences

In the case where there are 30 sequences, we try to build one or more consensus sequences. For that, we first cluster sequences according to their similarity score. We compute all pairwise alignments in semi-global mode. A pair of sequences is considered *compatible* if the alignement is above 85% identity. We also look at *included* sequences (a sequence is a substring of the other sequence). This allows us to build a *compatibility matrix*, containing either 0 (not compatible), 1 (compatible) or 2 (included) for each pair of alignments. We then use a greedy clustering approach to build groups of similar sequences, where all pairs of sequences are either of value 1 or 2. Starting by the most frequent non-included adapter, we add all included sequences to the cluster and then all sequences compatible with all previously selected sequences. For each cluster of size at least 2, we then contruct a multiple sequence alignment. One consensus sequence is computed per cluster using a simple vote for each nucleotide. Singleton sequences (that could not be clustered with any other sequences) are rejected.

Pairwise alignments and multiple sequence alignments are computed with the SeqAn library using the functions `globalAlignment` and `globalMsaAlignment` respectively.

#### 2.4.3 Poor consensus warning

At the end, each consensus sequence is a putative *adapter sequence* with an associated *frequency level*: the relative size of the cluster. When all frequency levels are below 30%, we raise a warning to express the fact that the prediction is not conformed to what is expected with untrimmed datasets.

### 2.5 Parameters

The algorithm depends on a series of parameters. They all have default values that work well in practice on a large variety of datasets (see section 4). Those parameters are as follows.

- Number of reads in each sample set (subsection 2.4): By default, the value is 40,000.
- Length of start and end regions selected for each read (subsection 2.1.1). By default, the value is 100.
- Length of the *k*-mers (subsection 2.1.1): By default, the value is 16.
- Number of top *k*-mers in the selection of frequent *k*-mers (subsection 2.1.1): by default, the value is 500.
- Low complexity threshold: The value depends on the *k*-mer length. By default, it equals 1 for *k*-mers of length 16. It is adjusted automatically for other values of *k* (subsection 2.1.2)

They can be customized by the user in the config file `ab_initio.config`

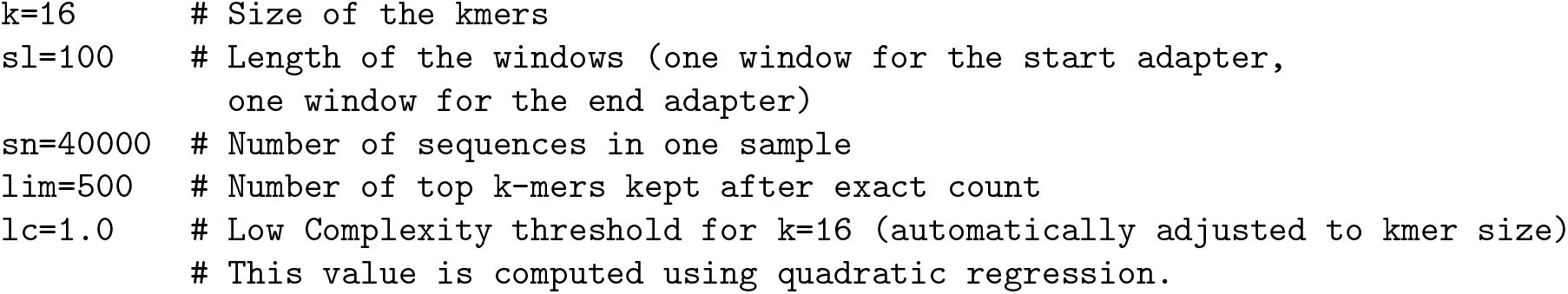

## 3 Implementation

The algorithm is implemented in C++ and Python, using the SEQAN library [6] and the NetworkX library^1^. The SEQAN library provides Optimal Search Schemes together with a bidrectional Burrows-Wheeler Transform and a FM-index for reads indexing, which makes the search of approximate *k*-mers very efficient [3, 2]. NetworkX is a graph library that is used in the assembly step of the algorithm.

This new code is available as an extension of Porechop to form a new software: Porechop_ABI. More precisely, the algorithm presented in Section 2 is implemented in the ABI module (*ab initio*), which is interfaced with Porechop. Porechop_ABI, as a whole, allows to automatically infer adapters and trim them in a single run. In practice, adapter sequences found by the ABI module are loaded in the database used by Porechop (file `adapter.py`). This organization is summarized in Figure 1. It is also possible for the user to only run the ABI module. In this specific case, the output is simply a set a putative start adapters and end adapters, when such sequences are extracted from the raw reads.

## 4 Experimental results

To assess the performances of Porechop_ABI, we selected a variety of datasets that correspond to the standard usage of the tool: ONT raw sequencing reads which have not been previously trimmed. We considered both simulated reads, and real DNA or cDNA reads generated with several flowcells, sequencing kits and base callers. The list is in Table 1. To complement this analysis, we also tested Porechop_ABI on a dataset trimmed with Guppy, which revealed that some residual fragments of adapters remain in the data.

**Table 1:** List of tested datasets

| Organism, tissue | Type | Flowcell | Sequencing Kit | Base caller | Source |
| --- | --- | --- | --- | --- | --- |
| Mouse | cDNA | Simulated with BadReads |  |  |  |
| *Eucalyptus pauciflora* | DNA | r9.5 | SQK-LSK108 | Albacore 2.0.2 | SRR7153074.1 |
| *Prunus dulcis* | DNA | r9.4 | SQK-NSK007 | Metrichor | SRA ERR3430401 |
| Mouse brain | cDNA | r9.4 | SQK-LSK008 | Metrichor | SRA PRJEB25574 |
| Human B-lymphocyte | cDNA | r9.4 | SQK-PCS108 | Albacore 2.1 |  |
| *Zea mays* | cDNA | r9.4 | SQK-PCS108 | Guppy | SRA PRJNA643165 |

For all of these datasets, the software was launched with default parameters, such as defined in the config file (see Subsection 2.5). The command-line is the following:

~~~
python3 ~/Porechop_ABI/porechop-runner.py -i $1 -go -abc $2 -v 0 >> results.txt
~~~

where $1 is the path to the tested dataset and $2 is the path to the config file.

All tests were ran on a HP-Compaq-Pro-6300-MT with an Intel(R) Core(TM) i5-3570 CPU @ 3.40GHz 16Go RAM with four threads. For each dataset, the additional processing time taken by Porechop_ABI compared to the original Porechop is less than 30 seconds per sample. Note that for this benchmark, we concentrate on the accuracy of the adapter sequences found. So the computation time does not include the trimming step, only the inference of the adapters (`--guess_adapter_only` option). In practice, both steps, inference and trimming, can be achieved with a single command:

~~~
porechop -abi -i input_file.fastq -o output_file.fastq
~~~

### 4.1 Simulated Nanopore reads

The first dataset is composed of 983,330 artificial long reads generated using BadRead (v0.1.5) [11] with default parameters. BadRead documentation specifies that the default adapters sequences inserted are Nanopore ligation adapters. Each adapter has a probability to appear in a given read (rate) and variable amount of the adapter will be visible (amount). The sequence for reads starts is AATGTACTTCGTTCAGTTACGTATTGCT (28nt) with a rate of 90% and an amount of 60%, the sequence for reads ends is GCAATACGTAACTGAACGAAGT (22nt) with a of rate 50% and an amount of 20% (more details are available in the BadRead README).

We used the GRCm38 assembly of the mouse transcriptome as a template to generate the dataset using the following command-line:

~~~
badread simulate --reference Mus_musculus.GRCm38.cdna.fasta
--quantity 2G >
simulated_badreads_1M.fastq
~~~

This reference transcriptome is available from ftp://ftp.ensembl.org/pub/release-102/fasta/mus_musculus/cdna/Mus_musculus.GRCm38.cdna.abinitio.fa.gz.

Figure 4 shows the results obtained with Porechop_ABI. Regarding the start adapter, a single sequence is found with 100% support. It is 27nt long, and matches the start adapter of BadReads. The only difference is that the first nucleotide is missing, and that the predicted sequence has an extra nucleotide `T` at the end. Regarding the end adapter, one again, there is a single sequence found, with 100% frequency. This a 21nt long sequence. The last nucleotide is missing compared to the Badreads end adapter. This result is very promising, because only half of the reads are intented to contain the end adapter (rate=50%), with a mean length of 20% of the original adapter across the whole dataset. Even in this case, Porechop_ABI is able to correctly recover a large part of the signal. In both cases, the difference between the predicted sequence and the reference sequence consists in a single extra of missing nucleotide at 5’ and 3’ ends, which means the sequence can be used for trimming.

**Figure 4:**
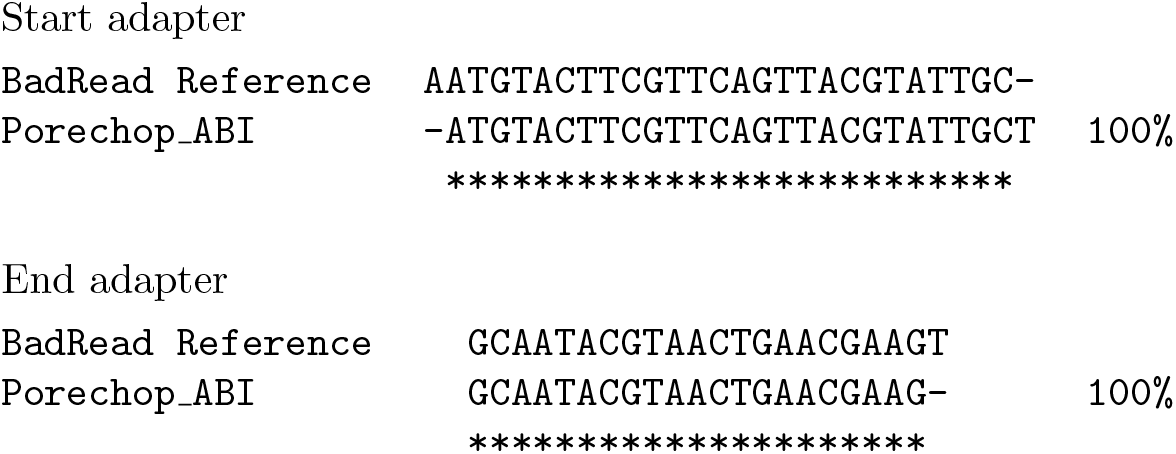
Adapters found by Porechop_ABI for simulated ONT reads (Section 4.1). The percentage indicated with the Porechop_ABI sequence is the support score, computed during the sampling phase of the algorithm. We mark identical positions between the reference and our sequence by stars.

### 4.2 Genome of Eucalyptus pauciflora

In [9], the authors generated high coverage of Nanopore reads (174×) from a single *E. pauciflora* individual. They prepared 1D ligation libraries according to the ONT’s protocol, SQK-LSK108 (whose adapters are identical to SQK-NSK007), and sequenced the reads using MinKNOW v1.7.3 with R9.5 flowcells on a MinION sequencer. They performed base calling with Albacore v2.0.2.

The whole project was deposited at NCBI under BioProject number PRJNA450887 and the whole-genome sequencing data are available in the SRA with accession number SRR7153044-SRR7153116. In this archive, we used dataset SRR7153074.1, that contains 818,267 reads with average read length 7959nt.

On this dataset, Porechop_ABI finds a single start adapter sequence and a single end adapter sequence, both with support 100%. Results are shown in Figure 5. As for the start region, adapter found by Porechop_ABI is very similar to the top adapter of SQK-NSK007-Y: SQK-NSK007-Y-top is 28 nt long, and we correctly recovered 26 nucleotides at the 3’ end. Our sequence does, however, contain 4 extranucleotides at the 5’ extremity. One can legitimately ask whether this difference have an impact on the search for the adapter in the sequences when trimming out the reads. To answer this question, we trimmed all reads of the dataset using the SQK-NSK007_Y adapters on the one hand, and the predicted adapters on the other hand, and compared the results. Out of the 818,267 reads, 393,174 reads are trimmed out with the SQK-NSK007_Y_top adapter (48.05% total), and 391,243 reads are trimmed with porechop_ABI adapters (47.81% total). Those numbers are very closed and, what is more, they refer to the same reads: 364,108 reads were trimmed out for both adapters and 397,958 reads had no adapters found by both tools. Regarding the end region, the majority of reads (74%) have not been trimmed out with either the SQK-NSK007_Y_bottom adapter or the adapter found with Porechop_ABI. For the remaining reads, they is a string overlap (more than 60%) between two methods. See Figure 6 for a Venn diagram.

**Figure 5:**
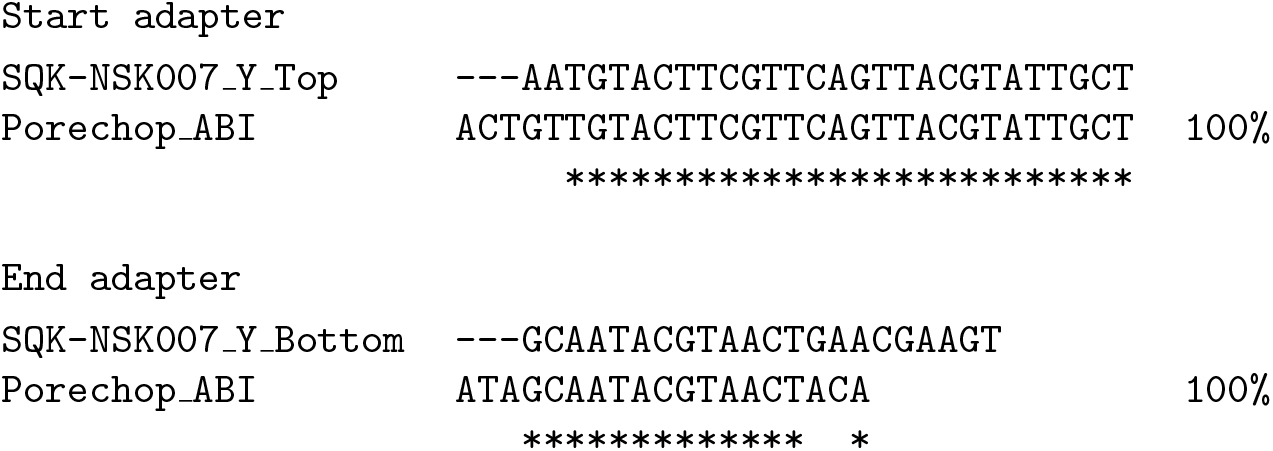
Adapters found by Porechop_ABI for the Eucalyptus reads (section 4.2). We align the sequences found against the SQK-NSK007_Y_Top and SQK-NSK007_Y_Bottom sequences, and mark identical positions with a star. The percentage indicated at the end of the Porechop_ABI line is the support score.

**Figure 6:**
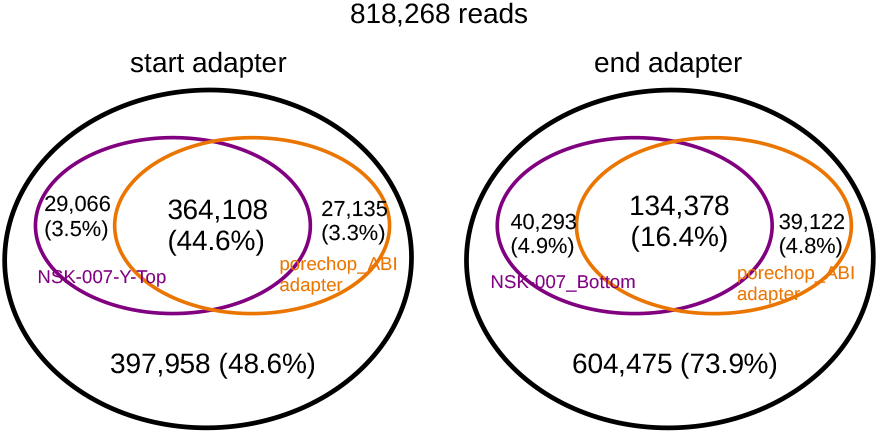
Venn diagram for Eucalyptus reads: we indicate the proportions of reads whoch have been identically trimmed out using the SQK-NSK007_Y adapter and our adapter.

#### Stability accross sampling

To go further with this first real dataset, we then tested *sampling stability* by running the program 100 times independently. For the start adapter, all runs produced exactly the same output, with 100% support at each try. For the end adapter, there are some minor variations: 91 tests obtained the same sequence with maximal support (100%), 8 tests obtained the same sequence with a lower support (98.3%) and one test produced a sequence with one extra nucleotide. See Table 2.

**Table 2:**
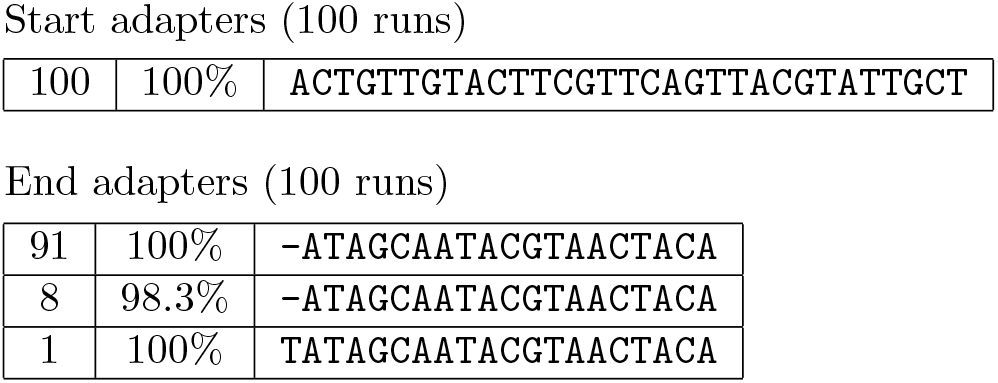
Stability across sampling(section 4.2). We ran the software 100 times, and for each run we report the adapter sequences found together with their support score. The first column is the number of runs, the second column is the support and the third column is the sequence. We obtained exactly the same start sequence for each of the 100 trials, with support 100%. As for the end adapter, 99 out of 100 runs returned the same sequence with support ranging from 98.3 to 100, and one run produced a sequence with one extra nucleotide at the beginning, with support 100.

### 4.3 Almond genome

This dataset comes from the sequencing of the *Prunus dulcis* genome accessible on SRA (ref: ERR3430401). It was sequenced on a r9.4 flowcell using the kit SQK-NSK007 and no prior PCR step [1]. It contains 441,203 reads.

Porechop_ABI finds one sequence for the start adapter, and one sequence for the end adapter both with 100% support. As shown in Figure 7, the start adapter matches closely to the SQK-NSK007_Y_Top adapter, as expected. The end adapter matches to the beginning of the SQK-NSK007_Y_Bottom adapter, meaning probably that most reads were cut before the end of the adapter.

**Figure 7:**
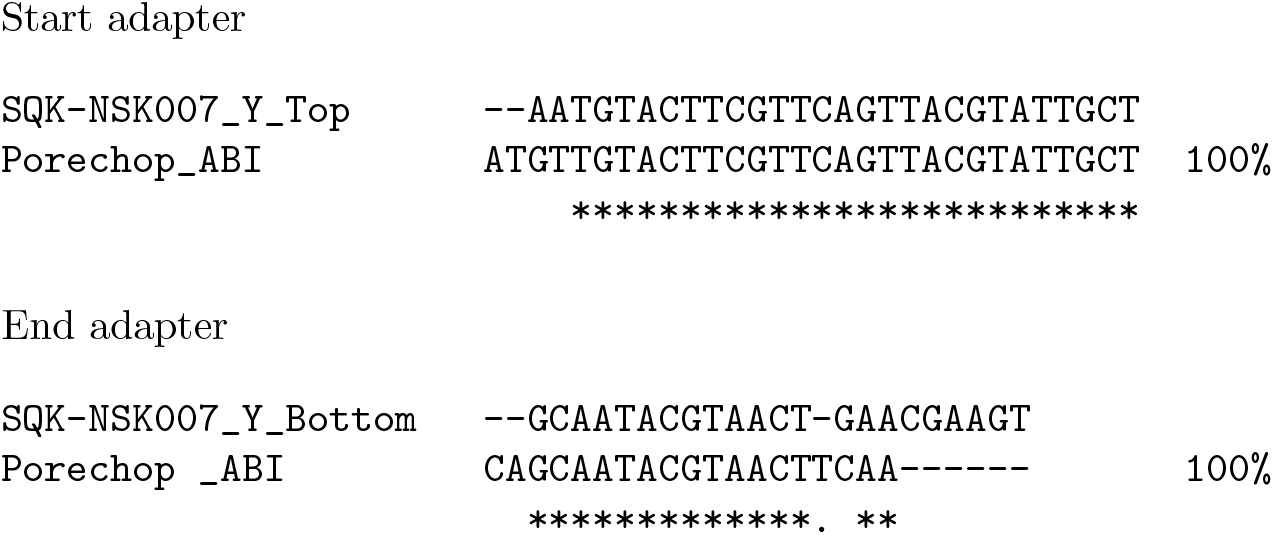
Start and end adapters found by Porechop_ABI for the almond genome dataset (section 4.3)

### 4.4 Mouse brain cDNA

This dataset contains 1D cDNA reads sequenced from mouse brain transcriptome using a Min-Ion equipped with a r9.4 flowcell. It was generated following a custom sequencing protocol: 2D cDNA synthesis (SQK-LSK008) followed by a 1D library protocol for sequencing.^2^. The dataset was basecalled using Metrichor 2.43.1 using 1D workflow 1D Basecalling RNN for LSK108, and contains 1,256,967 reads. It is publicly available on ENA/SRA with the id PRJEB25574 (filename: BYK_CB_ONT_1_FAF04998_A_1Donly).

In this dataset, Porechop_ABI identified two distinct sequences for the start region, both with support 50%. Those sequences have approximately the same length: 78nt for the first one, and 74 nt for the other one. They have the same initial motif and the same final motif. They differ with the middle part. When comparing all these parts to adapters present in the Porechop database `adapter.py`, it appears that each of them can be aligned to known adpater, as shown in Figure 8. The first part matches perfectly with SQK-NSK007_Y_Top, the last part is SQK-MAP006_Short_Y_Top_LI32, and the middle part corresponds respectively to the PCR_1_start and PCR_2_start. This organization with three adapters is consistent with ONT guidelines: ”*this kind of 3-motif pattern is apparently expected with this chemistry. The first part of your motif is the adapter, the middle part is the primer and the last part is the leader (the PCA adapter is y shaped and the rightmost portion of the sequence is the MAP006 short)*”^3^.

**Figure 8:**
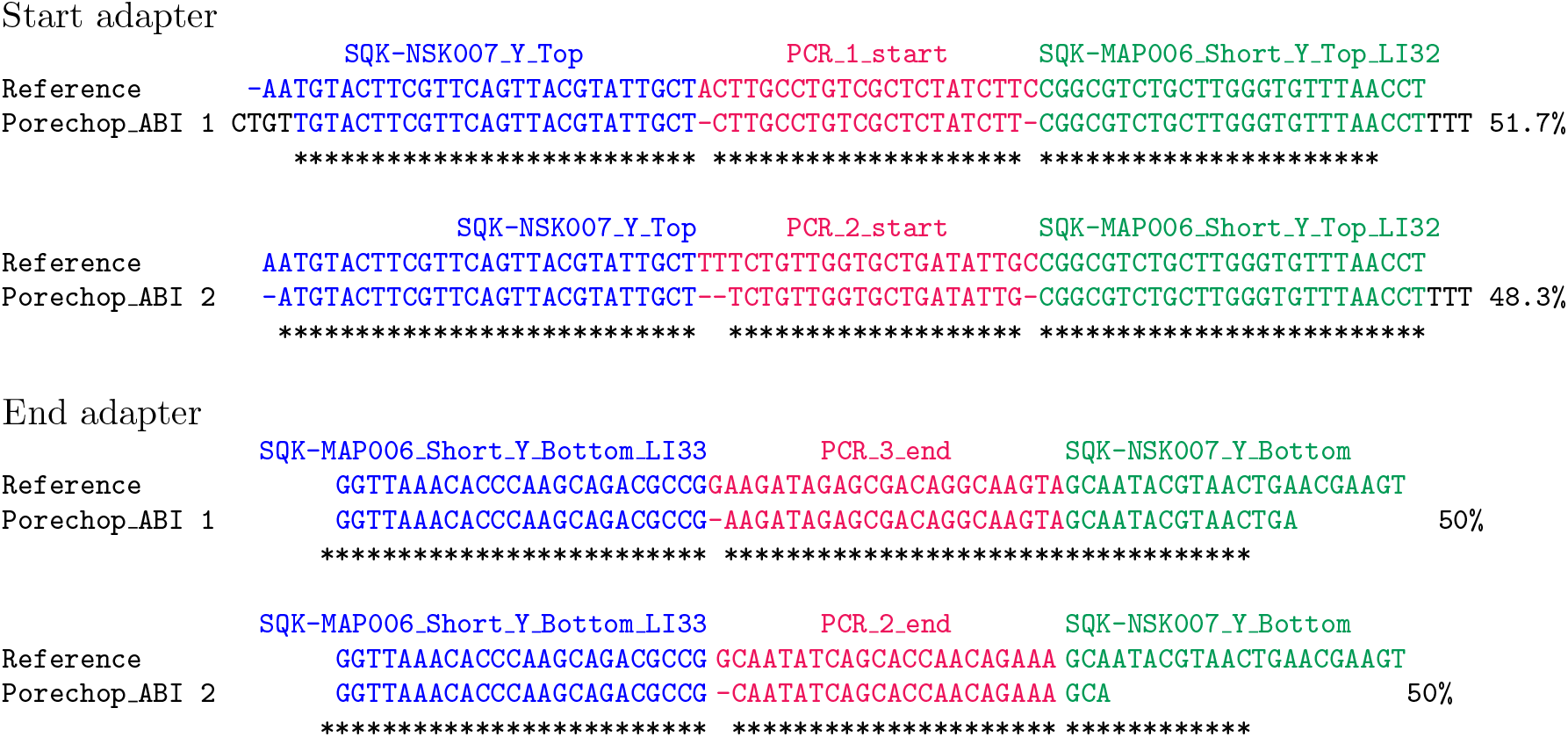
Start and end adapters for the mouse brain dataset (section 4.4)

Similarly, the program finds two sequences in the 3’ end region, with respective support 58.3% and 41.3%, and both sequences exhibit a 3-motif organization. The beginning part is SQK-NSK007_Y_Top, the end is SQK-MAP006_Short_Bottom_LI33, and the middle part corresponds to two distinct PCR adapters, PCR_2_end and PCR_3_end.

As a conclusion, this example shows the ability of Porechop_ABI to successfully manage datasets with mixed adapters. This distinctive feature of the tool comes from the clustering and consensus step in the core module, described in subsection 2.4.2.

### 4.5 Data from the Nanopore WGS consortium

The Nanopore WGS Consortium^4^ sequenced a human poly(A) transcriptome from B-lymphocyte cell line (GM12878) [12]. Two datasets from two different sequencing centers were selected for this benchmark. The first center is Bham (University of Birmingham, file `Bham_Run1_20171115_1D.pass.dedup.fastq`), and the second is UCSC (University of California, Santa Cruz, file `UCSC_Run1_20170919_1D.pass.dedup.fastq`). Both datasets were sequenced using the 1D cDNA protocol, a r9.4 flowcell and the SQK-PCS108 kit. They were basecalled using Albacore 2.1. Adapter sequences for the SQK-PCS108 kit are not documented or referenced in Porechop database. All we know is that the sequencing protocol involves a PCR step.

We ran Porechop_ABI on the two datasets independently, with the idea in mind that since experimental conditions are similar, we should obtain similar results from these two datasets. The results are presented in Figure 9. For the start adapter, we found two sequences with UCSC dataset, both with support 50%, and a single sequence with the BHAM dataset. All these three sequences begin with the same prefix:

**Figure 9:**
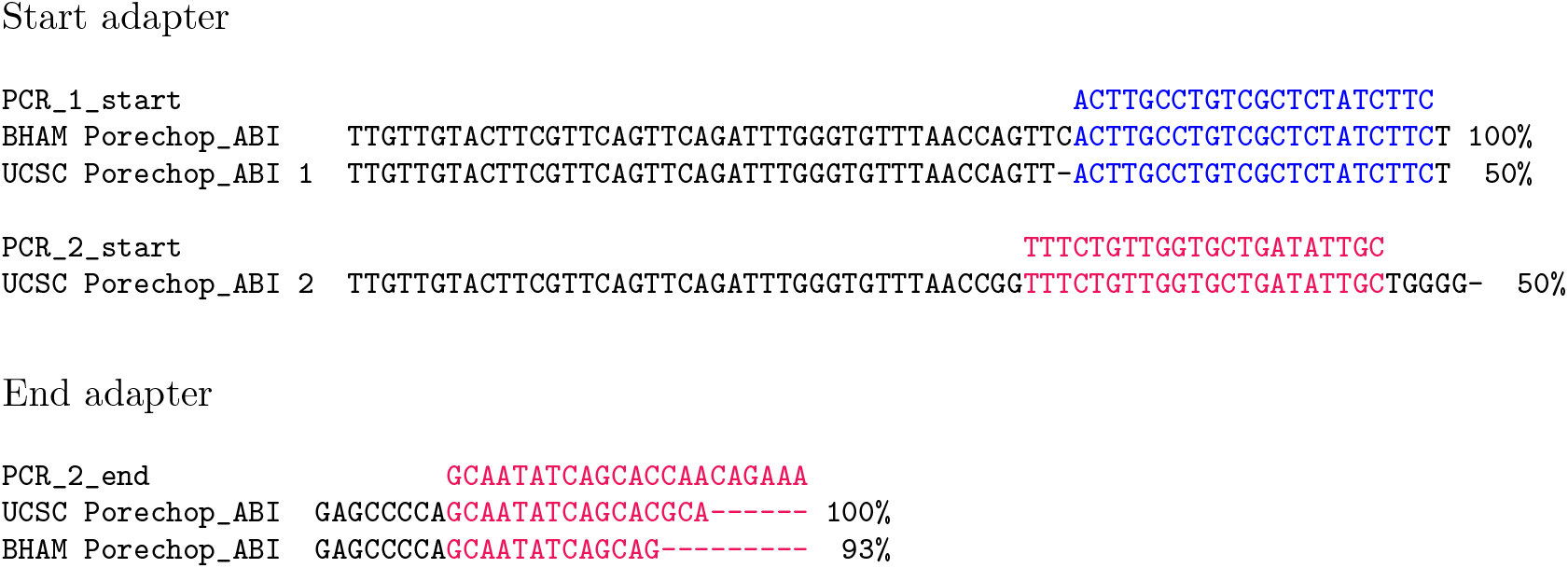
Porechop_ABI results for UCSC and BHAM human datasets

~~~
TTGTTGTACTTCGTTCAGTTCAGATTTGGGTGTTTAACCAG
~~~

Then adapter 1 found on UCSC dataset and the single adapter found on BHAM dataset both match to the PCR1 start adapter, while adapter 2 of UCSC dataset matches the PCR2 start adapter. The presence of PCR adapters is expected with the standard 1d cDNA protocol, since a PCR step is involved.

As for the end adapter, both datasets give a single sequence. The end adapters is much shorter, and only a partial alignment of the PCR_2_end adapter was possible.

### 4.6 Zea mays: testing basecaller trimming with Guppy

All previously processed datasets were basecalled with either Metrichor or Albacore, which both require a downstream trimming phase. In this example, we tested Porechop_ABI on a dataset basecalled with Guppy, that supposedly trims out adapters at the basecalling stage. Our goal is to test Guppy trimming ability, and check whether or not Porechop_ABI could detect leftover adapters.

We used a *Zea mays* 1D cDNA dataset picked from SRA: PRJNA643165. It was basecalled with Guppy with trimming mode activated. It contains a total of 5,396,492 reads obtained using a r9.4 flowcell and the SQK-PCS108 sequencing kit (like the human cDNA datasets of Section 4.5).

Results are shown on Figure 10. In both cases, start and end regions, Porechop_ABI found a stable adapter sequence. The start sequence closely matches the PCR_2_start adapter sequence with some excess on both sides. The analysis of the end adapter raises additional questions. The sequence maps partially on the PCR_2_end adapter. This confirms the fact that parts of the PCR_2 adapter sequences were still be found in this dataset. Since no other adapter sequences were discovered, we suggest that Guppy may have been able to trim out other adapter sequences.

**Figure 10:**
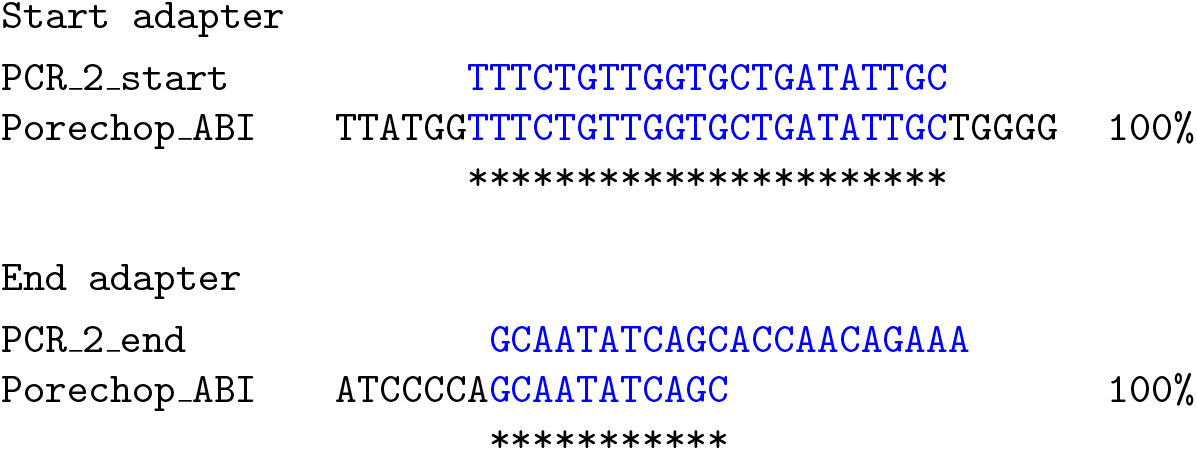
Adapters for the Guppy maize dataset (section 4.6)

In order to look at the prevalence of residual adaptor sequences, we used Porechop in trimming mode with our predicted adapters on the entire dataset (5,396,492 reads). The result is as follows:

- 1,699,193 reads contained start adapters in the 1-150 region, resulting in 103,923,858 base pairs removed.
- 2,102,217 reads contained end adapters from position −150, resulting in 38,466,512 base pairs removed.
- 2,257 reads were split based on middle adapters.

It means that 31% reads still contain traces of the start adapter near the 5’ extremity, and 39% reads traces of the end adapter near the 3’ extremity. The rate of middle adapters, found in the middle part, is approximately 4%. It is an indication that the dataset may contain some chimeric reads.

If this behavior is consistent on all datasets trimmed by Guppy, it is more than likely that some adapter sequences are still present in a lot of recent public dataset.

## 5 Negative controls

In this section, we provide negative controls, in order to evaluate the selectivity of the tool. We look at the existence of false negative predictions: what is the outcome of Porechop_ABI when there is no adapter sequence to find ?

### 5.1 Random sequences

The first test is made of 100 datasets composed of 1 million randomly generated sequences of length 100nt (independently distributed sequences with 25% A’s, 25% C’s, 25% G’s and 25% T’s). We ran Porechop_ABI on each of these datasets. It did not find any adapter sequence in all cases.

### 5.2 Control from the eucalytpus genome

A second negative control was built from the *E. pauciflora* dataset of Section 4.2. Since adapters are intended to be inserted at the extremities of the reads, alternative regions of the reads are supposed to be adapter-free, except for some chimera reads. To test whether Porechop_ABI would find motifs in the middle of the reads, we selected all reads whose length is greater than 700 nt and ran the algorithm on the region [500, 600]. We did this 100 times, in order to test the stability. For each of these 100 runs, Porechop_ABI issued a *low frequency warning*, meaning that the *k*-mers used to build the sequences are questionable (see subsection 2.3.3). In some cases, the program was able to build consensus sequences and output adapter sequences, but none of these sequences exceed the recommended 30% support threshold, and the poor consensus warning was activated in all cases (see subsection 2.4.3). It means that Poerchop ABI did not find any motif.

### 5.3 Control from the brain mouse transcriptome

The last negative control was built from the mouse brain cDNA dataset of Section 4.4, looking at windows where start and end adapters are not supposed to be found. The first window is [201, 300] and the second window is [−300, −201] in vicinity to the 3’ end of the reads. In both cases, the *low frequency warning* was triggered, but a mild signal was found. On the start area, Porechop_ABI finds a sequence of length 52nt, visible in Figure 11. A Blast search showed that this pattern exactly maps on the coding sequence of the myelin basic protein (NM_001025251.2), which is a major constituent of the myelin sheath of oligodendrocytes and Schwann cells in the nervous system. For the other window, we found two sequences, that are also visible on Figure 11. Blast search establishes that those two sequences are fragments of the myelin basic protein and of the carboxypeptidase E that is involved in the biosynthesis of most neuropeptides (NM_013494.4).

**Figure 11:**
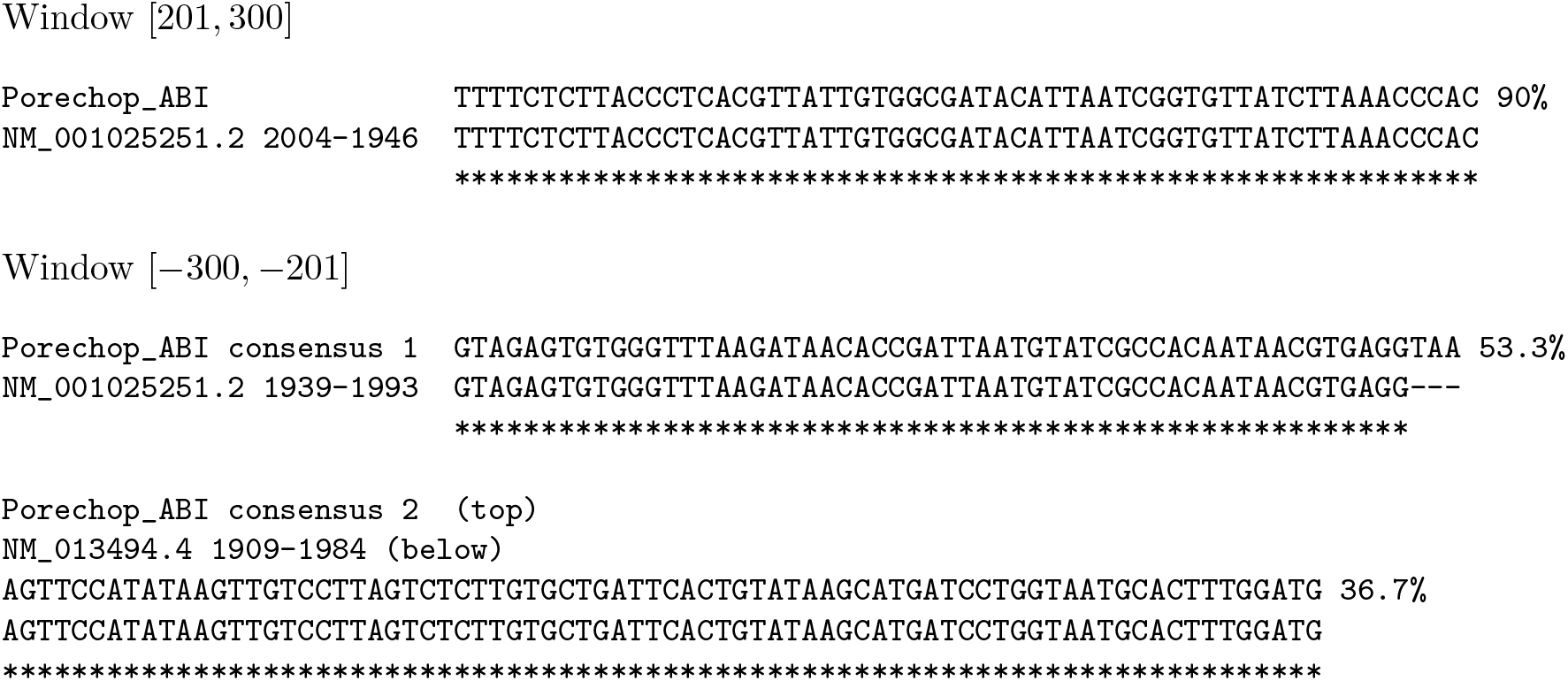
Adapters for negative control dataset built from mouse brain transcriptome (subsection 5.3). NM_001025251.2 is the transcript identifier for the *Mus musculus* myelin basic protein, and NM_013494.4 the transcript identifier for the *Mus musculus* carboxypeptidase E. For both windows, the total support is strictly smaller than 100%, because the method also found orphan sequences whose support is below the 30% threshold.

So in both windows, Porechop_ABI indicates that there is no adapter left, and finds fragments of genes that are known to be expressed in brain cells. This example shows that the tool is highly sensitive and is able detect a low biogical signal, even if this not its primary purpose. The warning messages are solid indications that there is no adapter present, and that the sequences found a must be treated with caution.

## 6 Discussion

We have developed a new software that meets the initial requirements: to infer the adapter sequences in ONT reads, without any prior knowledge about the adapters. It allows to determine whether the reads have already been trimmed out, and if not, which adapters have been used. This algorithm is integrated into the open source software Porechop, which makes it easy to use and allows trimming out in a single pass once the adapters have been identified.

We believe that Porechop_ABI can be useful to help analysing freshly sequenced data, by verifying that the reads indeed contain the expected adapters or that they have been accurately trimmed out. It also facilitates the usage of data available on public repositories, that often lack metadata on that matter.

## Footnotes

1 https://networkx.org/

2 Personal communication, Genoscope

3 ONT customer service, personal communication

4 https://github.com/nanopore-wgs-consortium/NA12878

